# Bayesian co-estimation of selfing rate and locus-specific mutation rates for a partially selfing population

**DOI:** 10.1101/020537

**Authors:** Benjamin D. Redelings, Seiji Kumagai, Liuyang Wang, Andrey Tatarenkov, Ann K. Sakai, Stephen G. Weller, Theresa M. Culley, John C. Avise, Marcy K. Uyenoyama

**Affiliations:** Department of Biology, Box 90338, Duke University, Durham, NC 27708-0338; Department of Ecology and Evolutionary Biology, University of California, Irvine, Irvine, CA 92697-2525; Department of Biological Sciences, University of Cincinnati, Cincinnati, OH 45220

**Keywords:** selfing rate, Ewens Sampling Formula, Bayesian, MCMC, mating system

## Abstract

We present a Bayesian method for characterizing the mating system of populations reproducing through a mixture of self-fertilization and random outcrossing. Our method uses patterns of genetic variation across the genome as a basis for inference about pure hermaphroditism, androdioecy, and gynodioecy. We extend the standard coalescence model to accommodate these mating systems, accounting explicitly for multilocus identity disequilibrium, inbreeding depression, and variation in fertility among mating types. We incorporate the Ewens Sampling Formula (ESF) under the infinite-alleles model of mutation to obtain a novel expression for the likelihood of mating system parameters. Our Markov chain Monte Carlo (MCMC) algorithm assigns locus-specific mutation rates, drawn from a common mutation rate distribution that is itself estimated from the data using a Dirichlet Process Prior model. Among the parameters jointly inferred are the population-wide rate of self-fertilization, locus-specific mutation rates, and the number of generations since the most recent outcrossing event for each sampled individual.

---

Inbreeding has pervasive consequences throughout the genome, affecting genealogical relationships between genes held at each locus within individuals and among multiple loci. This generation of genome-wide, multilocus disequilibria of various orders transforms the context in which evolution proceeds. Here, we address a simple form of inbreeding: a mixture of self-fertilization (selfing) and random outcrossing (Clegg 1980; Ritland 2002).

Various methods exist for the estimation of selfing rates from genetic data. Wright’s (1921) fundamental approach bases the estimation of selfing rates on the coefficient of inbreeding (*F*_*IS*_), which reflects the departure from Hardy-Weinberg proportions of genotypes for a given set of allele frequencies. The maximum likelihood method of Enjalbert and David (2000) detects inbreeding from departures of multiple loci from Hardy-Weinberg proportions, estimating allele frequencies for each locus and accounting for correlations in heterozygosity among loci (identity disequilibrium, Cockerham and Weir 1968). David *et al.* (2007) extend the approach of Enjalbert and David (2000), basing the estimation of selfing rates on the distribution of heterozygotes across multiple, unlinked loci, while accommodating errors in scoring heterozygotes as homozygotes. A primary objective of 

~~~
InStruct
~~~

 (Gao *et al.* 2007) is the estimation of admixture. It extends the widely-used program 

~~~
structure
~~~

 (Pritchard *et al.* 2000), which bases the estimation of admixture on disequilibria of various forms, by accounting for disequilibria due to selfing. Progeny array methods (see Ritland 2002), which base the estimation of selfing rates on the genetic analysis of progeny for which one or more parents are known, are particularly well-suited to plant populations. Wang *et al.* (2012) extend this approach to a random sample of individuals by reconstructing sibship relationships within the sample.

Methods that base the estimation of inbreeding rates on the observed departure from random union of gametes require information on expected Hardy-Weinberg proportions. Population-wide frequencies of alleles observed in a sample at locus *l* ({*p*_*li*_}) can be estimated jointly in a maximum-likelihood framework (*e.g.*, Hill *et al.* 1995) or integrated out as nuisance parameters in a Bayesian framework (*e.g.*, Ayres and Balding 1998). Similarly, locus-specific heterozygosity

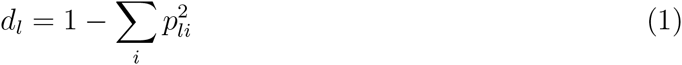

can be obtained from observed allele frequencies (Enjalbert and David 2000) or estimated directly and jointly with the selfing rate (David *et al.* 2007).

In contrast, our Bayesian method for the analysis of partial self-fertilization derives from a coalescence model that accounts for genetic variation and uses the Ewens Sampling Formula (ESF, Ewens 1972). Our approach replaces the estimation of allele frequencies or heterozygosity (1) by the estimation of a locus-specific mutation rate (*θ\**) under the infinite-alleles model of mutation. We use a Dirichlet Process Prior (DPP) to determine the number of classes of mutation rates, the mutation rate for each class, and the class membership of each locus. We assign the DPP parameters in a conservative manner so that it creates a new mutational class only if sufficient evidence exists to justify doing so. Further, while other methods assume that the frequency in the population of an allelic class not observed in the sample is zero, the ESF provides the probability, under the infinite-alleles model of mutation, that the next-sampled gene represents a novel allele (see (22a)).

To estimate the probability that a random individual is uniparental (*s\**), we exploit identity disequilibrium (Cockerham and Weir 1968), the correlation in heterozygosity across loci. This association, even among unlinked loci, reflects that all loci within an individual share a history of inbreeding back to the most recent random outcrossing event. Conditional on the number of generations since this event, the genealogical histories of unlinked loci are independent. Our method infers the number of consecutive generations of self-fertilization in the immediate ancestry of each sampled diploid individual and the probability of coalescence during this period between the lineages at each locus.

In inferring the full likelihood from the observed frequency spectrum of diploid genotypes at multiple unlinked loci, we determine the distributions of the allele frequency spectra ancestral to the sample at the most recent point at which all sampled gene lineages at each locus reside in separate individuals. At this point, the ESF provides the exact likelihood, obviating the need for further genealogical reconstruction. This approach permits computationally efficient analysis of samples comprising large numbers of individuals and large numbers of loci observed across the genome.

Here, we address the estimation of inbreeding rates in populations undergoing pure hermaphroditism, androdioecy (hermaphrodites and males), or gynodioecy (hermaphrodites and females). Our method provides a means for the simultaneous inference of various aspects of the mating system, including the population proportions of sexual forms and levels of inbreeding depression. We apply our method to simulated data sets to demonstrate its accuracy in parameter estimation and in assessing uncertainty. Our application to microsatellite data from the androdioecious killifish *Kryptolebias marmoratus* (Mackiewicz *et al.* 2006; Tatarenkov *et al.* 2012) and to the gynodioecious Hawaiian endemic *Schiedea salicaria* (Wallace *et al.* 2011) illustrates the formation of inferences about a number of biologically significant aspects, including measures of effective population size.

## Evolutionary model

We describe our use of the Ewens Sampling Formula (ESF, Ewens 1972) to determine likelihoods based on a sample of diploid multilocus genotypes.

From a reduced sample, formed by subsampling a single gene from each locus from each diploid individual, one could use the ESF to determine a likelihood function with a single parameter: the mutation rate, appropriately scaled to account for the acceleration of the coalescence rate caused by inbreeding (Nordborg and Donnelly 1997; Fu 1997). Observation of diploid genotypes provides information about another parameter: the probability that a random individual is uniparental (uniparental proportion). We describe the dependence of these two composite parameters on the basic parameters of models of pure hermaphroditism, androdioecy, and gynodioecy.

### Rates of coalescence and mutation

Here, we describe the structure of the coalescence process shared by our models of pure hermaphroditism, androdioecy, and gynodioecy.

#### Relative rates of coalescence and mutation

We represent the probability that a random individual is uniparental by *s\** and the probability that a pair of genes that reside in distinct individuals descend from the same parent in the immediately preceding generation by 1*/N \**. These quantities determine the coalescence rate and the scaled mutation rate of the ESF.

A pair of lineages residing in distinct individuals derive from a single parent (P) in the preceding generation at rate 1*/N \**. They descend from the same gene (immediate coalescence) or from distinct genes in that individual with equal probability. In the latter case, P is either uniparental (probability *s\**), implying descent once again of the lineages from a single individual in the preceding generation, or biparental, implying descent from distinct individuals. Residence of a pair of lineages in a single individual rapidly resolves either to coalescence, with probability

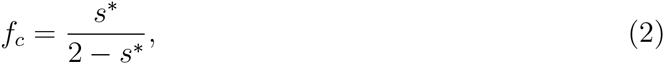

or to residence in distinct individuals, with the complement probability. This expression is identical to the classical coefficient of identity (Wright 1921; Haldane 1924). The total rate of coalescence of lineages sampled from distinct individuals corresponds to

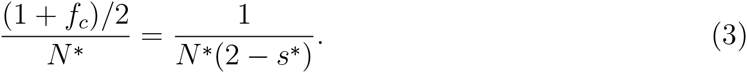

Our model assumes that coalescence and mutation occur on comparable time scales:

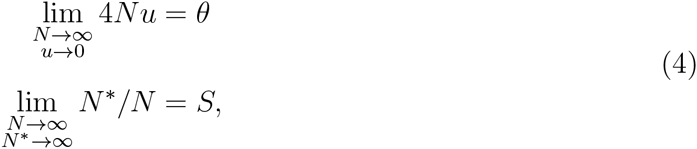

for *u* the rate of mutation under the infinite alleles model and *N* an arbitrary quantity that goes to infinity at a rate comparable to *N \** and 1*/u*. Here, *S* represents a scaled measure of effective population size (termed “inbreeding effective size” by Crow and Denniston 1988), relative to a population comprising *N* reproductives.

In large populations, switching of lineages between uniparental and biparental carriers occurs on the order of generations, virtually instantaneously relative to the rate at which lineages residing in distinct individuals coalesce (Nordborg and Donnelly 1997; Fu 1997). Our model assumes independence between the processes of coalescence and mutation and that these processes occur on a much longer time scale than random outcrossing:

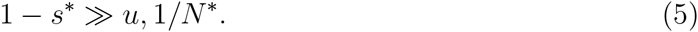

For *m* lineages, each residing in a distinct individual, the probability that the most recent event corresponds to mutation is

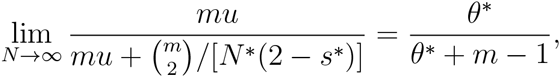

in which

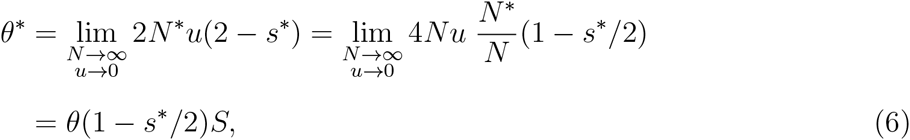

for *θ* and *S* defined in (4). In inbred populations, the single parameter of the ESF corresponds to *θ\**.

#### Uniparental proportion and the rate of parent-sharing

In a population comprising *N*_*h*_ hermaphrodites, the rate of parent-sharing corresponds to 1*/N*_*h*_, and the uniparental proportion (*s\**) corresponds to

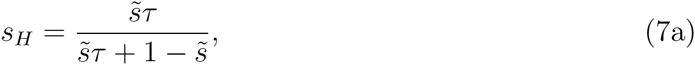

for 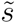 the fraction of uniparental offspring at conception and *τ* the rate of survival of uniparental relative to biparental offspring. For the pure-hermaphroditism model, we assign the arbitrary constant *N* in (4) as *N*_*h*_, implying

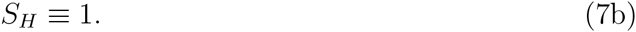

In androdioecious populations, comprising *N*_*h*_ reproducing hermaphrodites and *N*_*m*_ reproducing males (female-steriles), the uniparental proportion (*s\**) is identical to the case of pure hermaphroditism (7)

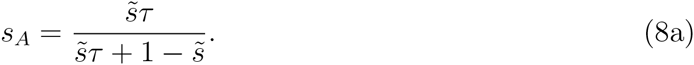

A random gene derives from a male in the preceding generation with probability

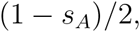

and from a hermaphrodite with the complement probability. A pair of genes sampled from distinct individuals derive from the same parent (1*/N \**) with probability

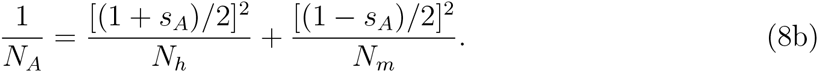

In the absence of inbreeding (*s*_*A*_ = 0), this expression agrees with the classical harmonic mean expression for effective population size (Wright 1969). For the androdioecy model, we assign the arbitrary constant in (4) as the number of reproductives (*N*_*h*_ + *N*_*m*_), implying a scaled rate of coalescence corresponding to

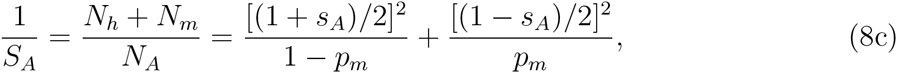

for

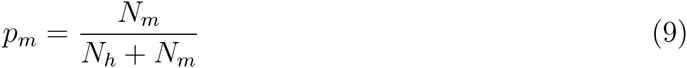

the proportion of males among reproductive individuals. Relative effective number *S*_*A*_ *∈*(0, 1] takes its maximum for populations in which the effective number *N*_*A*_, implied by the rate of parent sharing, corresponds to the total number of reproductives (*N*_*A*_ = *N*_*h*_ +*N*_*m*_). At *S*_*A*_ = 1, the probability that a random gene derives from a male parent equals the proportion of males among reproductives:

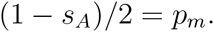

In gynodioecious populations, in which *N*_*h*_ hermaphrodites and *N*_*f*_ females (male-steriles) reproduce, the uniparental proportion (*s\**) corresponds to

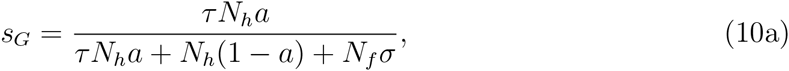

in which *σ* represents the seed fertility of females relative to hermaphrodites and *a* the proportion of seeds of hermaphrodites set by self-pollen. A random gene derives from a female in the preceding generation with probability

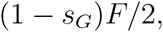

for

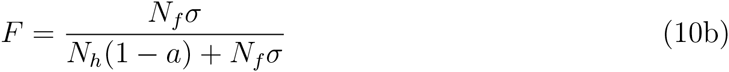

the proportion of biparental offspring that have a female parent. A pair of genes sampled from distinct individuals derive from the same parent (1*/N \**) with probability

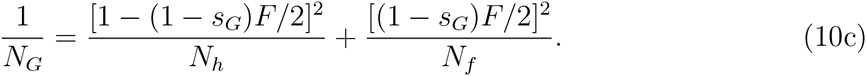

We assign the arbitrary constant *N* in (4) as (*N*_*h*_ + *N*_*f*_), implying a scaled rate of coalescence of

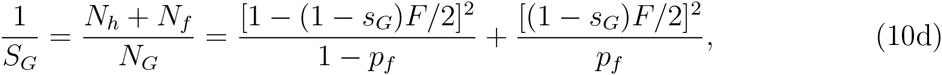

for

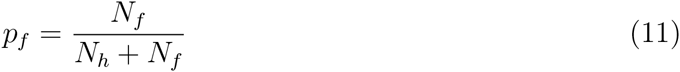

the proportion of females among reproductive individuals. As for the androdioecy model, *S*_*G*_ *∈* (0, 1] achieves its maximum only if the proportion of females among reproductives equals the probability that a random gene derives from a female parent:

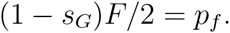

### Likelihood

We here address the probability of a sample of diploid multilocus genotypes.

#### Genealogical histories

For a sample comprising up to two alleles at each of *L* autosomal loci in *n* diploid individuals, we represent the observed genotypes by

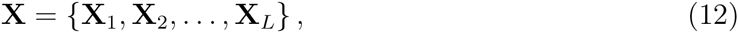

in which **X**_*l*_ denotes the set of genotypes observed at locus *l*,

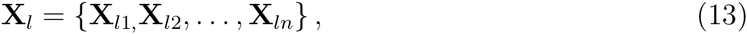

with

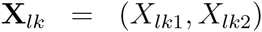

the genotype at locus *l* of individual *k*, with alleles *X*_*lk*1_ and *X*_*lk*2_.

To facilitate accounting for the shared recent history of genes borne by an individual in sample, we introduce latent variables

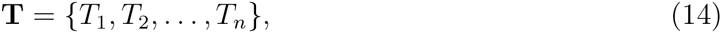

for *T*_*k*_ denoting the number of consecutive generations of selfing in the immediate ancestry of the *k*^*th*^ individual, and

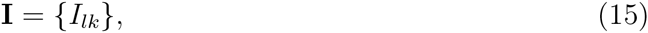

for *I*_*lk*_ indicating whether the lineages borne by the *k*^*th*^ individual at locus *l* coalesce within the most recent *T*_*k*_ generations. Independent of other individuals, the number of consecutive generations of inbreeding in the ancestry of the *k*^*th*^ individual is geometrically distributed:

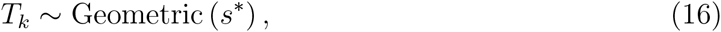

with *T*_*k*_ = 0 signifying that individual *k* is the product of random outcrossing. Irrespective of whether 0, 1, or 2 of the genes at locus *l* in individual *k* are observed, *I*_*lk*_ indicates whether the two genes at that locus in individual *k* coalesce during the *T*_*k*_ consecutive generations of inbreeding in its immediate ancestry:

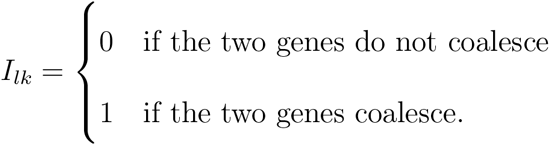

Because the pair of lineages at any locus coalesce with probability 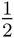 in each generation of selfing,

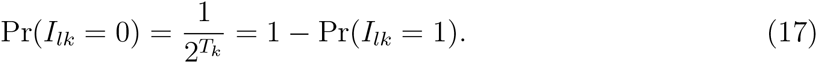

Figure 1 depicts the recent genealogical history at a locus *l* in 5 individuals. Individuals 2 and 5 are products of random outcrossing (*T*_2_ = *T*_5_ = 0), while the others derive from some positive number of consecutive generations of selfing in their immediate ancestry (*T*_1_ = 2, *T*_3_ = 3, *T*_4_ = 1). Both individuals 1 and 3 are homozygotes (*αα*), with the lineages of individual 3 but not 1 coalescing more recently than the most recent outcrossing event (*I*_*l*1_ = 0, *I*_*l*3_ = 1). As individual 2 is heterozygous (*αβ*), its lineages necessarily remain distinct since the most recent outcrossing event (*I*_*l*2_ = 0). One gene in each of individuals 4 and 5 are unobserved (*\**), with the unobserved lineage in individual 4 but not 5 coalescing more recently than the most recent outcrossing event (*I*_*l*4_ = 1, *I*_*l*5_ = 0).

**Figure 1.**
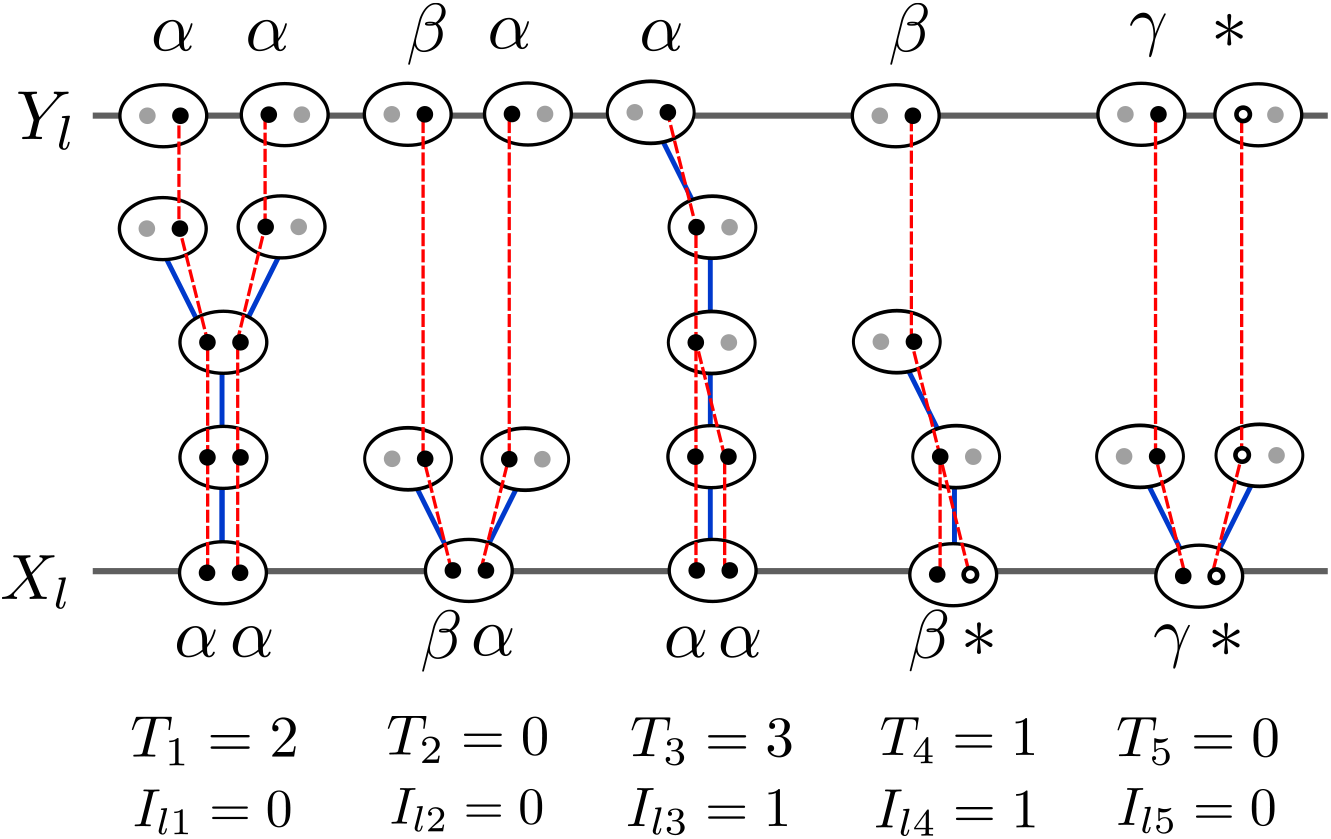
Following the history of the sample (**X**_*l*_) backwards in time until all ancestors of sampled genes reside in different individuals (**Y**_*l*_). Ovals represent individuals and dots represent genes. Blue lines indicate the parents of individuals, while red lines represent the ancestry of genes. Filled dots represent sampled genes for which the allelic class is observed (Greek letters) and their ancestral lineages. Open dots represent genes in the sample with unobserved allelic class (*). Grey dots represent other genes carried by ancestors of the sampled individuals. The relationship between the observed sample **X**_*l*_ and the an-cestral sample **Y**_*l*_ is determined by the intervening coalescence events **I**_*l*_. **T** indicates the number of consecutive generations of selfing for each sampled individual.

In addition to the observed sample of diploid individuals, we consider the state of the sampled lineages at the most recent generation in which an outcrossing event has occurred in the ancestry of all *n* individuals. This point in the history of the sample occurs 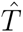 generations into the past, for

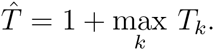

In Figure 1, for example, 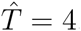, reflecting the most recent outcrossing event in the ancestry of individual 3. The ESF provides the probability of the allele frequency spectrum at this point.

We represent the ordered list of allelic states of the lineages at 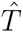 generations into the past by

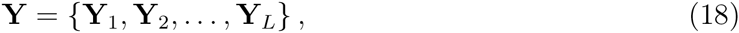

for **Y**_*l*_ a list of ancestral genes in the same order as their descendants in **X**_*l*_. Each gene in **Y**_*l*_ is the ancestor of either 1 or 2 genes at locus *l* from a particular individual in **X**_*l*_ (13), depending on whether the lineages held by that individual coalesce during the consecutive generations of inbreeding in its immediate ancestry. We represent the number of genes in **Y**_*l*_ by *m*_*l*_ (*n ≤ m*_*l*_ *≤* 2*n*). In Figure 1, for example, **X**_*l*_ contains 10 genes in 5 individuals, but **Y**_*l*_ contains only 8 genes, with *Y*_*l*1_ the ancestor of only the first allele of **X**_*l*1_ and *Y*_*l*5_ the ancestor of both alleles of **X**_*l*3_.

We assume (5) that the initial phase of consecutive generations of selfing is sufficiently short to ensure a negligible probability of mutation in any lineage at any locus and a negligible probability of coalescence between lineages held by distinct individuals more recently than 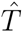. Accordingly, the coalescence history **I** (15) completely determines the correspondence between genetic lineages in **X** (12) and **Y** (18).

#### Computing the likelihood

In principle, the likelihood of the observed data can be computed from the augmented likelihood by summation:

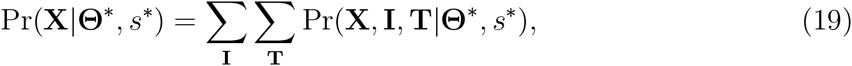

for

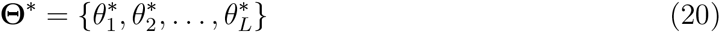

the list of scaled, locus-specific mutation rates, *s\** the population-wide uniparental proportion for the reproductive system under consideration (*e.g.*, (7) for the pure hermaphroditism model), and **T** (14) and **I** (15) the lists of latent variables representing the time since the most recent outcrossing event and whether the two lineages borne by a sampled individual coalesce during this period. Here we follow a common abuse of notation in using Pr(**X**) to denote Pr(**X** = **x**) for random variable **X** and realized value **x**. Summation (19) is computationally expensive: the number of consecutive generations of inbreeding in the immediate ancestry of an individual (*T*_*k*_) has no upper limit (compare David *et al.* 2007) and the number of combinations of coalescence states (*I*_*lk*_) across the *L* loci and *n* individuals increases exponentially (2^*Ln*^) with the total number of assignments. We perform Markov chain Monte Carlo (MCMC) to avoid both these sums.

To calculate the augmented likelihood, we begin by applying Bayes rule:

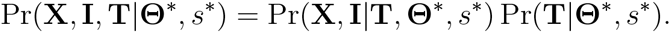

Because the times since the most recent outcrossing event **T** depend only on the uniparental proportion *s\**, through (16), and not on the rates of mutation **θ\***,

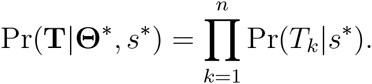

Even though our model assumes the absence of physical linkage among any of the loci, the genetic data **X** and coalescence events **I** are not independent across loci because they depend on the times since the most recent outcrossing event **T**. Given **T**, however, the genetic data and coalescence events are independent across loci

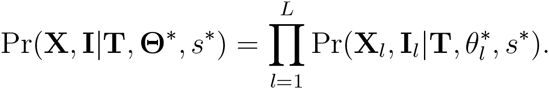

Further,

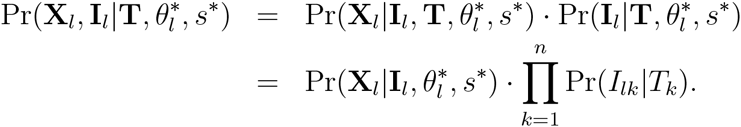

This expression reflects that the times to the most recent outcrossing event **T** affect the observed genotypes **X**_*l*_ only through the coalescence states **I**_*l*_ and that the coalescence states **I**_*l*_ depend only on the times to the most recent outcrossing event **T**, through (17).

To compute 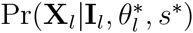, we incorporate latent variable **Y**_*l*_ (18), describing the states of lineages at the most recent point at which all occur in distinct individuals (Figure 1):

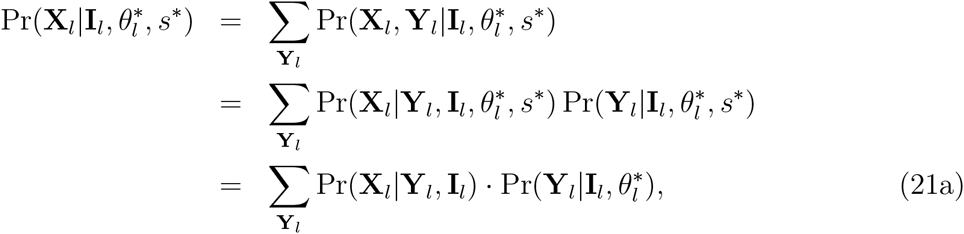

reflecting that the coalescence states **I**_*l*_ establish the correspondence between the spectrum of genotypes in **X**_*l*_ and the spectrum of alleles in **Y**_*l*_ and that the distribution of **Y**_*l*_, given by the ESF, depends on the uniparental proportion *s\** only through the scaled mutation rate 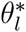 (6).

Given the sampled genotypes **X**_*l*_ and coalescence states **I**_*l*_, at most one ordered list of alleles **Y**_*l*_ produces positive Pr(**X**_*l*_|**Y**_*l*_, **I**_*l*_) in (21a). Coalescence of the lineages at locus *l* in any heterozygous individual (*e.g.*, *X*_*lk*_ = (*β, α*) with *I*_*lk*_ = 1 in Figure 1) implies

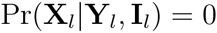

for all **Y**_*l*_. Any non-zero Pr(**X**_*l*_|**Y**_*l*_, **I**_*l*_) precludes coalescence in any heterozygous individual and **Y**_*l*_ must specify the observed alleles of **X**_*l*_ in the order of observation, with either 1 (*I*_*lk*_ = 1) or 2 (*I*_*lk*_ = 0) instances of the allele for any homozygous individual (*e.g.*, *X*_*lk*_ = (*α, α*)). For all cases with non-zero Pr(**X**_*l*_|**Y**_*l*_, **I**_*l*_),

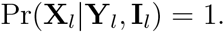

Accordingly, expression (21a) reduces to

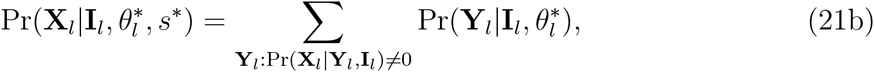

a sum with either 0 or 1 terms. Because all genes in **Y**_*l*_ reside in distinct individuals, we obtain 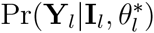 from the Ewens Sampling Formula for a sample, of size

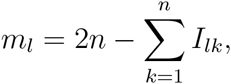

ordered in the sequence in which the genes are observed.

To determine 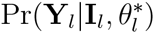 in (21b), we use a fundamental property of the ESF (Ewens 1972; Karlin and McGregor 1972): the probability that the next-sampled (*i*^*th*^) gene represents a novel allele corresponds to

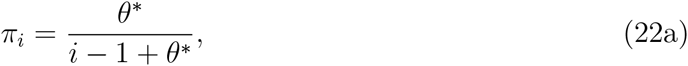

for *θ\** defined in (6), and the probability that it represents an additional copy of already-observed allele *j* is

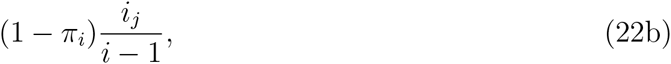

for *i*_*j*_ the number of replicates of allele *j* in the sample at size 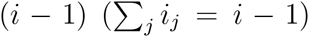. Appendix A presents a first-principles derivation of (22a). Expressions (22) imply that for **Y**_*l*_ the list of alleles at locus *l* in order of observance,

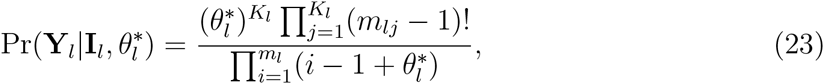

in which *K*_*l*_ denotes the total number of distinct allelic classes, *m*_*lj*_ the number of replicates of the *j*^*th*^ allele in the sample, and *m*_*l*_ =∑_*j*_ *m*_*lj*_ the number of lineages remaining at time 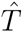 (Figure 1).

#### Missing data

Our method allows the allelic class of one or both genes at each locus to be missing. In Figure 1, for example, the genotype of individual 4 is **X**_*l*4_ = (*β, \**), indicating that the allelic class of the first gene is observed to be *β*, but that of the second gene is unknown.

A missing allelic specification in the sample of genotypes **X**_*l*_ leads to a missing specification for the corresponding gene in **Y**_*l*_ unless the genetic lineage coalesces, in the interval between **X**_*l*_ and **Y**_*l*_, with a lineage ancestral to a gene for which the allelic type was observed. Figure 1 illustrates such a coalescence event in the case of individual 4. In contrast, the lineages ancestral to the genes carried by individual 5 fail to coalescence more recently than their separation into distinct individuals, giving rise to a missing specification in **Y**_*l*_.

The probability of **Y**_*l*_ can be computed by simply summing over all possible values for each missing specification. Equivalently, those elements may simply be dropped from **Y**_*l*_before computing the probability via the ESF, the procedure implemented in our method.

## BAyesian inference framework

### Prior on mutation rates

Ewens (1972) showed for the panmictic case that the number of distinct allelic classes observed at a locus (*e.g.*, *K*_*l*_ in (23)) provides a sufficient statistic for the estimation of the scaled mutation rate. Because each locus *l* provides relatively little information about the scaled mutation rate 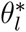 (6), we assume that mutation rates across loci cluster in a finite number of groups. However, we do not know *a priori* the group assignment of loci or even the number of distinct rate classes among the observed loci. We make use of the Dirichlet process prior to estimate simultaneously the number of groups, the value of *θ \** for each group, and the assignment of loci to groups.

The Dirichlet process comprises a base distribution, which here represents the distribution of the scaled mutation rate *θ\** across groups, and a concentration parameter *α*, which controls the probability that each successive locus forms a new group. We assign 0.1 to *α* of the Dirichlet process, and place a gamma distribution (γ(*α* = 0.25, *β* = 2)) on the mean scaled mutation rate for each group. As this prior has a high variance relative to the mean (0.5), it is relatively uninformative about *θ\**.

### Model-specific parameters

Derivations presented in the preceding section indicate that the probability of a sample of diploid genotypes under the infinite alleles model depends on only the uniparental proportion *s\** and the scaled mutation rates **θ\*** (20) across loci. These composite parameters are determined by the set of basic demographic parameters **Ψ** associated with each model of reproduction under consideration. As the genotypic data provide equal support to any combination of basic parameters that implies the same values of *s\** and **θ\***, the full set of basic parameters for any model are in general non-identifiable using the observed genotype frequency spectrum alone.

Even so, our MCMC implementation updates the basic parameters directly, with likelihoods determined from the implied values of *s\** and **θ\***. This feature facilitates the incorporation of information in addition to the genotypic data that can contribute to the estimation of the basic parameters under a particular model or assessment of alternative models. We have

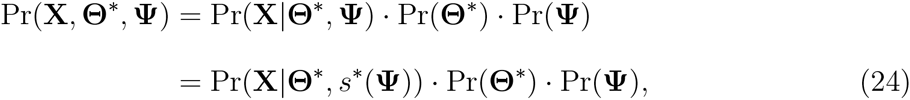

for **X** the genotypic data and *s\**(**Ψ**) the uniparental proportion determined by **Ψ** for the model under consideration. To determine the marginal distribution of *θ*_*l*_ (4) for each locus *l*, we use (6), incorporating the distributions of *s\**(**Ψ**) and *S*(**Ψ**), the scaling factor defined in (4):

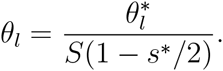

For the pure hermaphroditism model (7), 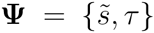, where 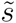 is the proportion of conceptions through selfing, and *τ* is the relative viability of uniparental offspring. We propose uniform priors for 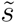 and *τ*:

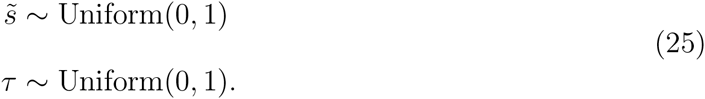

For the androdioecy model (8), we propose uniform priors for each basic parameter in 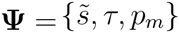

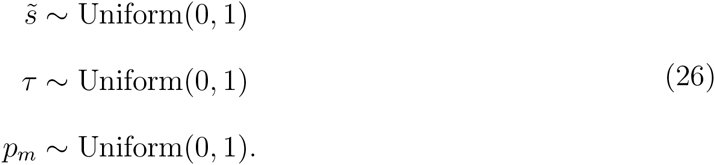

For the gynodioecy model (10), **Ψ** = {*a, τ, p*_*f*_, *σ*}, including *a* the proportion of egg cells produced by hermaphrodites fertilized by selfing, *p*_*f*_ (11) the proportion of females (male-steriles) among reproductives, and *σ* the fertility of females relative to hermaphrodites. We propose the uniform priors

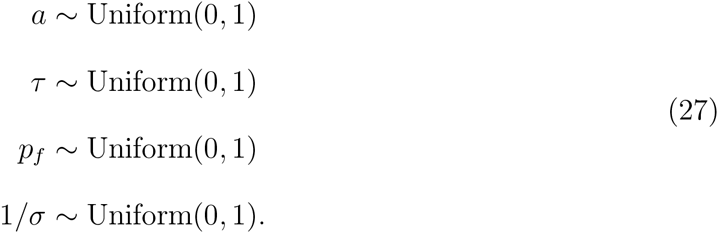

## Assessment of accuracy and coverage using simulated data

We developed a forward-in-time simulator (

~~~
https://github.com/skumagai/selfingsim
~~~

) that tracks multiple neutral loci with locus-specific scaled mutation rates (**θ**) in a population comprising *N* reproducing hermaphrodites of which a proportion *s\** are of uniparental origin. We used this simulator to generate data under two sampling regimes: large (*L* = 32 loci in each of *n* = 70 diploid individuals) and small (*L* = 6 loci in each of *n* = 10 diploid individuals). We applied our Bayesian method and 

~~~
RMES
~~~

 (David *et al.* 2007) to simulated data sets. A description of the procedures used to assess the accuracy and coverage properties of the three methods is included in the Supplementary Online Material.

In addition, we determine the uniparental proportion (*s\**) inferred from the departure from Hardy-Weinberg expectation (*F*_*IS*_, Wright 1969) alone. Our *F*_*IS*_-based estimate entails setting the observed value of *F*_*IS*_ equal to its classical expectation *s*/*(2 *- s\**) (Wright 1921; Haldane 1924) and solving for *s\**:

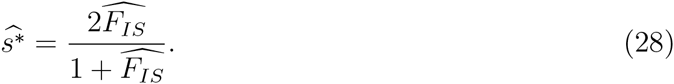

In accommodating multiple loci, this estimate incorporates a multilocus estimate for 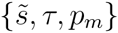 ---(Appendix B) but, unlike those generated by our Bayesian method and 

~~~
RMES
~~~

, does not use identity disequilibrium across loci within individuals to infer the number of generations since the most recent outcross event in their ancestry. As our primary purpose in examining the *F*_*IS*_-based estimate (28) is to provide a baseline for the results of those likelihood-based methods, we have not attempted to develop an index of error or uncertainty for it.

### Accuracy

To assess relative accuracy of estimates of the uniparental proportion *s\**, we determine the bias and root-mean-squared error of the three methods by averaging over 10^4^ data sets (10^2^ independent samples from each of 10^2^ independent simulations for each assigned *s\**). In contrast with the point estimates of *s\** produced by 

~~~
RMES
~~~

, our Bayesian method generates a posterior distribution. To facilitate comparison, we reduce our estimate to a single value, the median of the posterior distribution of *s\**, with the caveat that the mode and mean may show different qualitative behavior (see Supplementary Online Material).

Figure 2 indicates that both 

~~~
RMES
~~~

 and our method show positive bias upon application to data sets for which the true uniparental proportion *s\** is close to zero and negative bias for *s\** close to unity. This trend reflects that both methods yield estimates of *s\** constrained to lie between 0 and 1. In contrast, the *F*_*IS*_-based estimate (28) underestimates *s\** throughout the range, even near *s\** = 0 (*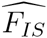* is not constrained to be positive). Our method has a bias near 0 that is substantially larger than the bias of 

~~~
RMES
~~~

, and an error that is slightly larger. A major contributor to this trend is that our Bayesian estimate is represented by only the median of the posterior distribution of the uniparental proportion *s\**. Figure 3 indicates that for data sets generated under a true value of *s\** of 0 (full random outcrossing), the posterior distribution for *s\** has greater mass near 0. Further, as the posterior mode does not display large bias near 0 (Figure S1), we conclude that the bias shown by the median (Figure 2) merely represents uncertainty in the posterior distribution for *s\** and not any preference for incorrect values. We note that our method assumes that the data are derived from a population reproducing through a mixture of self-fertilization and random outcrossing. Assessment of a model of complete random mating (*s\** = 0) against the present model (*s* >* 0) might be conducted through the Bayes factor.

**Figure 2.**
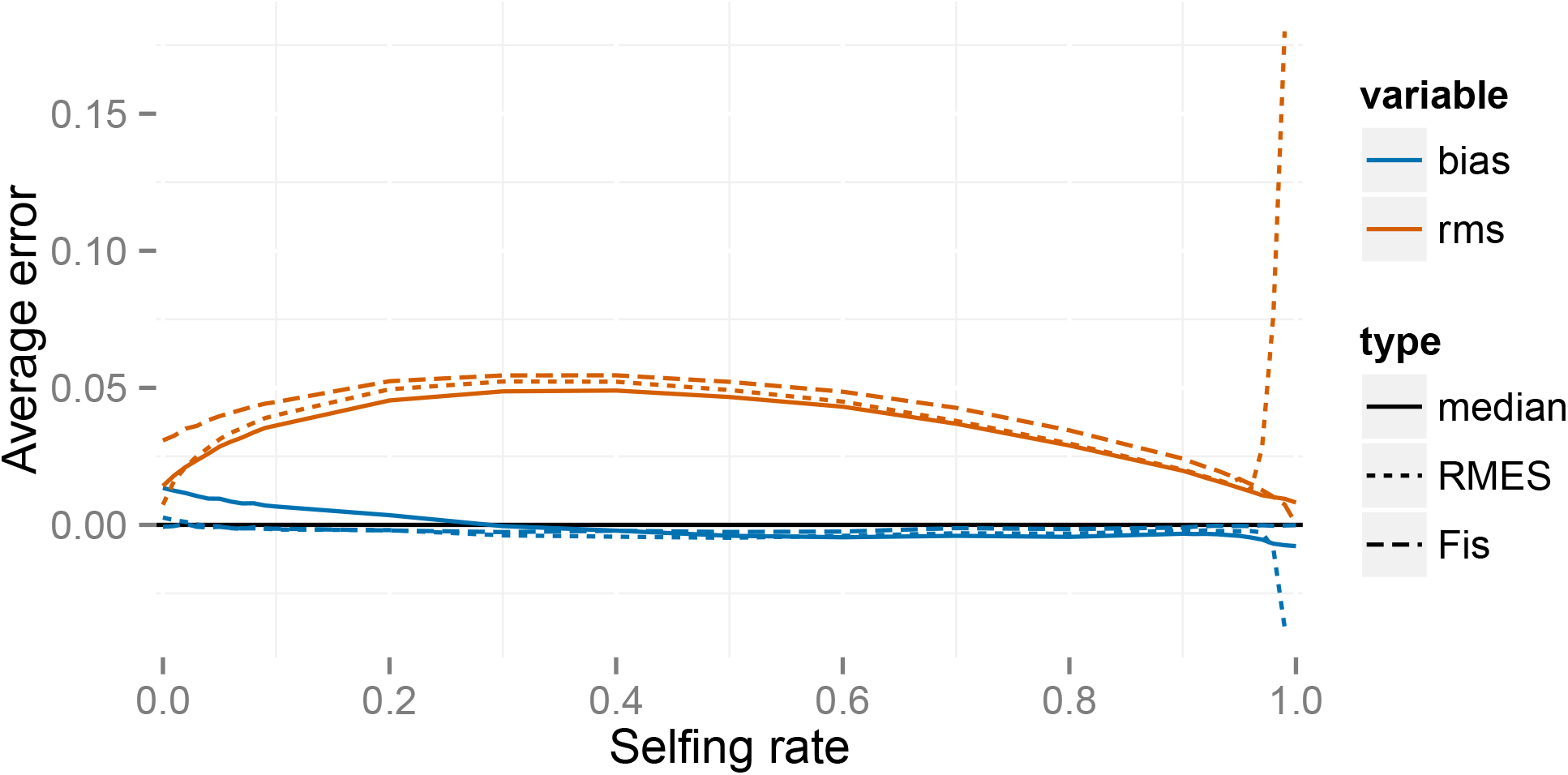
Errors for the full likelihood (posterior median), 

~~~
RMES
~~~

, and *F*_*IS*_-based (28) methods for a large simulated sample (*n* = 70 individuals, *L* = 32 loci). In the legend, rms indicates the root-mean-squared error and bias the average deviation. Averages are taken across simulated data sets at each true value of *s\**.

Except in cases in which the true *s\** is very close to 0, the error for 

~~~
RMES
~~~

 exceeds the error for our method under both sampling regimes (Figure 2). 

~~~
RMES
~~~

 differs from the other two methods in the steep rise in both bias and rms error for high values of *s\**, with the change point occurring at lower values of the uniparental proportion *s\** for the small sampling regime (*n* = 10, *L* = 6). A likely contributing factor to the increased error shown by 

~~~
RMES
~~~

 under high values of *s\** is its default assumption that the number of generations in the ancestry of any individual does not exceed 20. Violations of this assumption arise more often under high values of *s\**, possibly promoting underestimation of the uniparental proportion. Further, 

~~~
RMES
~~~

 discards data at loci at which no heterozygotes are observed, and terminates analysis altogether if the number of loci drops below 2. 

~~~
RMES
~~~

 treats all loci with zero heterozygosity (1) as uninformative, even if multiple alleles are observed. In contrast, our full likelihood method uses data from all loci, with polymorphic loci in the absence of heterozygotes providing strong evidence of high rates of selfing (rather than low rates of mutation). Under the large sampling regime (*n* = 70, *L* = 32), 

~~~
RMES
~~~

 discards on average 50% of the loci for true *s\** values exceeding 0.94, with less than 10% of data sets unanalyzable (fewer than 2 informative loci) even at *s\** = 0.99 (Figure 4). Under the *n* = 10, *L* = 6 regime, 

~~~
RMES
~~~

 discards on average 50% of loci for true *s\** values exceeding 0.85, with about 50% of data sets unanalyzable under *s* ≥* 0.94.

**Figure 4.**
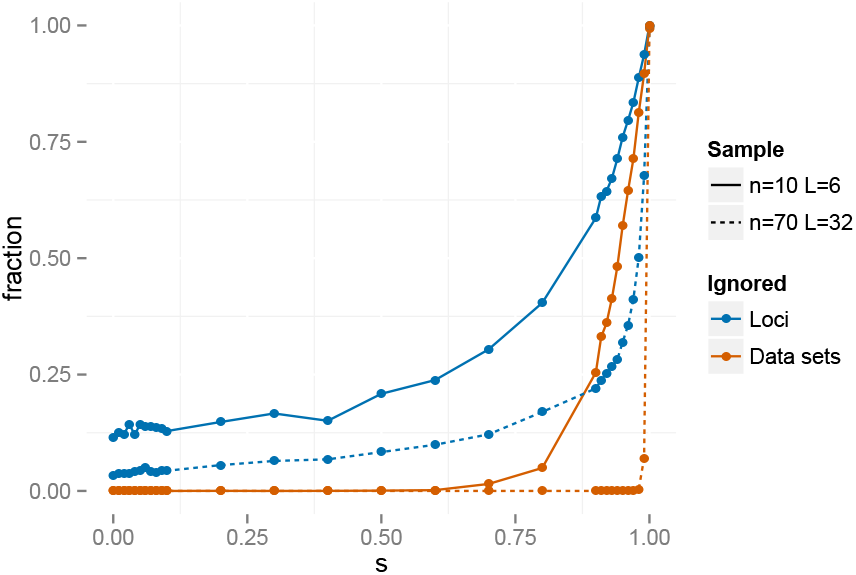
Fraction of loci and data sets that are ignored by 

~~~
RMES
~~~

.

The error for the *F*_*IS*_-based estimate (28) also exceeds the error for our method. It is largest near *s\** = 0 and vanishes as *s\** approaches 1, a pattern distinct from 

~~~
RMES
~~~

 (Figure 2).

### Coverage

We determine the fraction of data sets for which the confidence interval (CI) generated by 

~~~
RMES
~~~

 and the Bayesian credible interval (BCI) generated by our method contains the true value of the uniparental proportion *s\**. This measure of coverage is a frequentist notion, as it treats each true value of *s\** separately. A 95% CI should contain the truth 95% of the time for each specific value of *s\**. However, a 95% BCI is not expected to have 95% coverage at each value of *s\**, but rather 95% coverage averaged over values of *s\** sampled from the prior. Of the various ways to determine a BCI for a given posterior distribution, we choose to report the highest posterior density BCI (rather than the central BCI, for example).

Figure 5 indicates that coverage of the 95% CIs produced by 

~~~
RMES
~~~

 are consistently lower than 95% across all true *s\** values under the large sampling regime (*n* = 70 *L* = 32). Coverage appears to decline as *s\** increases, dropping from 86% for *s\** = 0.1 to 64% for *s\** = 0.99. In contrast, the 95% BCIs have slightly greater than 95% frequentist coverage for each value of *s\**, except for *s\** values very close to the extremes (0 and 1). Under very high rates of inbreeding (*s* ≈* 1), an assumption (5) of our underlying model (random outcrossing occurs on a time scale much shorter than the time scales of mutation and coalescence) is likely violated. We observed similar behavior under nominal coverage levels ranging from 0.5 to 0.99 (Supplementary Material).

**Figure 5.**
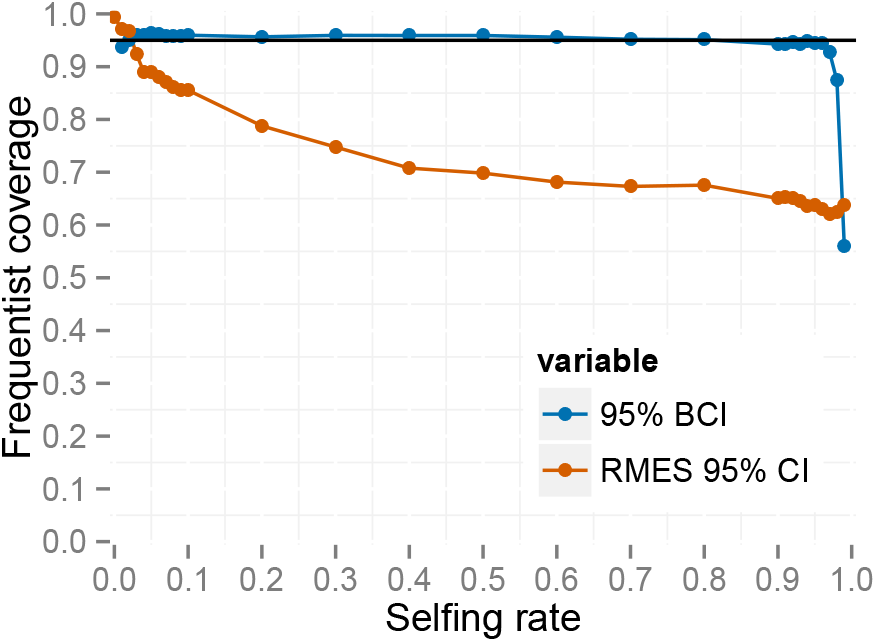
Frequentist coverage at each level of *s\** for 95% intervals from 

~~~
RMES
~~~

 and the method based on the full likelihood under the large sampling regime (*n* = 70, *L* = 32). 

~~~
RMES
~~~

 intervals are 95% confidence intervals computed via profile likelihood. Full likelihood intervals are 95% highest posterior density Bayesian credible intervals.

### Number of consecutive generations of selfing

In order to check the accuracy of our reconstructed generations of selfing, we examine the posterior distributions of selfing times *{T*_*k*_*}* for *s\** = 0.5 under the large sampling regime (*n* = 70, *L* = 32). We average posterior distributions for selfing times across 100 simulated data sets, and across individuals *k* = 1 *…* 70 within each simulated data set. We then compare these averages based on the simulated data with the exact distribution of selfing times across individuals (Figure 6). The pooled posterior distribution closely matches the exact distribution. This simple check suggests that our method correctly infers the true posterior distribution of selfing times for each sampled individual.

**Figure 6.**
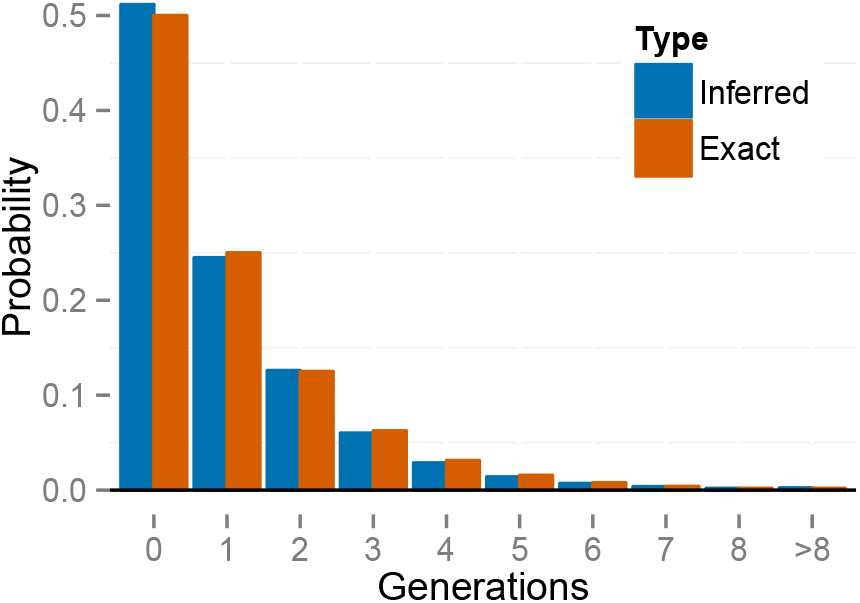
Exact distribution of selfing times under *s\** = 0.5 compared to the posterior distribution averaged across individuals and across data sets.

## Analysis of microsatellite data from natural populations

### Androdioecious vertebrate

Our analysis of data from the androdioecious killifish *Kryptolebias marmoratus* (Mackiewicz *et al.* 2006; Tatarenkov *et al.* 2012) incorporates genotypes from 32 microsatellite loci as well as information on the observed fraction of males. Our method simultaneously estimates the proportion of males in the population (*p*_*m*_) together with rates of locus-specific mutation (*θ\**) and the uniparental proportion (*s*_*A*_). We apply the method to two populations, which show highly divergent rates of inbreeding.

#### Parameter estimation

Our androdioecy model (25) comprises 3 basic parameters, including the fraction of males among reproductives (*p*_*m*_) and the relative viability of uniparental offspring (*τ*). Our analysis incorporates the observation of *n*_*m*_ males among *n*_*total*_ zygotes directly into the likelihood expression:

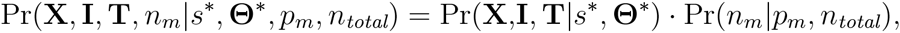

in which

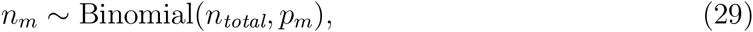

reflecting that *s\** and **Θ\*** are sufficient to account for **X**, **I**, and **T**, and also independent of *n*_*m*_, *n*_*total*_, and *p*_*m*_.

In the absence of direct information regarding the existence or intensity of inbreeding depression, we impose the constraint *τ* = 1 to permit estimation of the uniparental proportion *s*_*A*_ under a uniform prior:

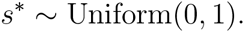

#### Low outcrossing rate

We applied our method to the BP data set described by Tatarenkov *et al.* (2012). This data set comprises a total of 70 individuals, collected in 2007, 2010, and 2011 from the Big Pine location on the Florida Keys.

Tatarenkov *et al.* (2012) report 21 males among the 201 individuals collected from various locations in the Florida Keys during this period, consistent with other estimates of about 1% (*e.g.*, Turner *et al.* 1992). Based on the long-term experience of the Tatarenkov–Avise laboratory with this species, we assumed observation of *n*_*m*_ = 20 males out of *n*_*total*_ = 2000 individuals in (29). We estimate that the fraction of males in the population (*p*_*m*_) has a posterior median of 0.01 with a 95% Bayesian Credible Interval (BCI) of (0.0062, 0.015).

Our estimates of mutation rates (*θ\**) indicate substantial variation among loci, with the median ranging over an order of magnitude (ca. 0.5–5.0) (Figure S4, Supplementary Material). The distribution of mutation rates across loci appears to be multimodal, with many loci having a relatively low rate and some having larger rates.

Figure 7 shows the posterior distribution of uniparental proportion *s*_*A*_, with a median of 0.95 and a 95% BCI of (0.93, 0.97). This estimate is somewhat lower than *F*_*IS*_-based estimate (28) of 0.97, and slightly higher than the 

~~~
RMES
~~~

 estimate of 0.94, which has a 95% Confidence Interval (CI) of (0.91, 0.96). We note that 

~~~
RMES
~~~

 discarded from the analysis 9 loci (out of 32) which showed no heterozygosity, even though 7 of the 9 were polymorphic in the sample.

**Figure 7.**
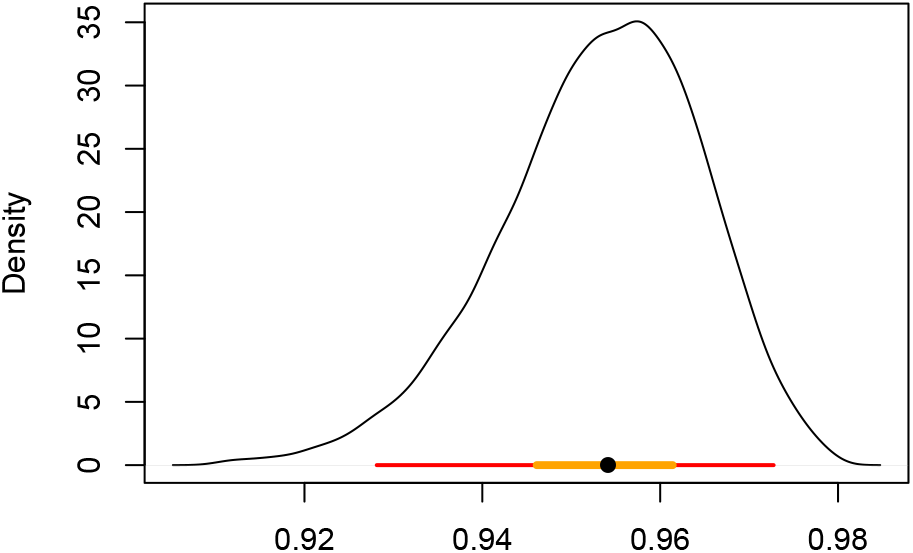
Posterior distribution of the uniparental proportion *s*_*A*_ for the BP population. The median is indicated by a black dot, with a red bar for the 95% BCI and an orange bar for the 50% BCI.

Our method estimates the latent variables *{T*_1_, *T*_2_, *…, T*_*n*_*}* (14), representing the number of generations since the most recent outcross event in the ancestry of each individual (Figure S5). Figure 8 shows the empirical distribution of the time since outcrossing across individuals, averaged over posterior uncertainty, indicating a complete absence of biparental individuals (0 generations of selfing). Because we expect that a sample of size 70 would include at least some biparental individuals under the inferred uniparental proportion (*s*_*A*_ *≈* 0.95), this finding suggests that any biparental individuals in the sample show lower heterozygosity than expected from the observed level of genetic variation. This deficiency suggests that an extended model that accommodates biparental inbreeding or population subdivision may account for the data better than the present model, which allows only selfing and random outcrossing.

**Figure 8.**
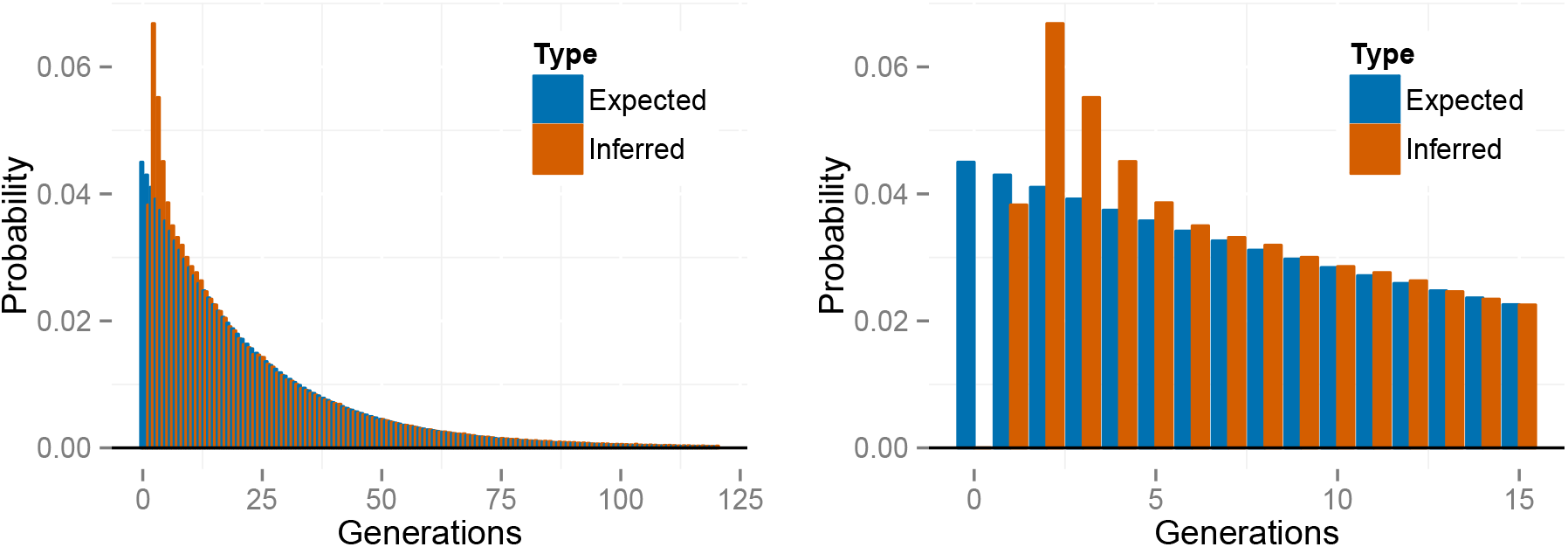
Empirical distribution of number of generations since the most recent outcross event (*T*) across individuals for the *K. marmoratus* (BP population), averaged across posterior samples. The right panel is constructed by zooming in on the panel on the left. “Expected” probabilities represent the proportion of individuals with the indicated number of selfing generations expected under the estimated uniparental proportion *s*_*A*_. “Inferred” probabilities represent proportions inferred across individuals in the sample. The first in-ferred bar with positive probability corresponds to *T* = 1.

#### Higher outcrossing rate

We apply the three methods to the sample collected in 2005 from Twin Cays, Belize (TC05: Mackiewicz *et al.* 2006). This data set departs sharply from that of the BP population, showing considerably higher incidence of males and levels of polymorphism and heterozygosity.

We incorporate the observation of 19 males among the 112 individuals collected from Belize in 2005 (Mackiewicz *et al.* 2006) into the likelihood (see (29)). Our estimate of the fraction of males in the population (*p*_*m*_) has a posterior median of 0.17 with a 95% BCI of (0.11, 0.25).

Figure S6 (Supplementary Material) indicates that the posterior medians of the locus-specific mutation rates range over a wide range (ca. 0.5–23). Two loci appear to exhibit a mutation rates substantially higher than other loci, both of which appear to have high rates in the BP population as well (Figure S4).

All three methods confirm the inference of Mackiewicz *et al.* (2006) of much lower in-breeding in the TC population relative to the BP population. Our posterior distribution of uniparental proportion *s*_*A*_ has a median and 95% BCI of 0.35 (0.25, 0.45) (Figure 9). median again lies between the *F*_*IS*_-based estimate (28) of 0:39 and the 

~~~
RMES
~~~

 estimate of 0:33, with its 95% CI of (0:30; 0:36). In this case, 

~~~
RMES
~~~

 excluded from the analysis only a single locus, which was monomorphic in the sample.

**Figure 9.**
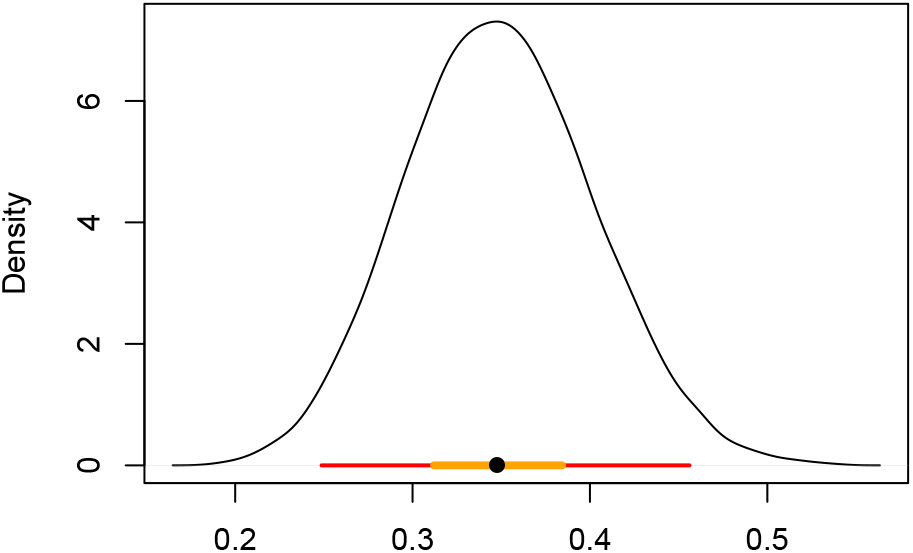
Posterior distribution of the uniparental proportion *s*_*A*_ for the TC population. Also shown are the 95% BCI (red), 50% BCI (orange), and median (black dot).

Figure 10 shows the inferred distribution of the number of generations since the most recent outcross event (*T*) across individuals, averaged over posterior uncertainty. In contrast to the BP population, the distribution of selfing time in the TC population appears to conform to the distribution expected under the inferred uniparental proportion (*s*_*A*_), including a high fraction of biparental individuals (*T*_*k*_ = 0). Figure S7 (Supplementary Material) presents the posterior distribution of the number of consecutive generations of selfing in the immediate ancestry of each individual.

**Figure 10.**
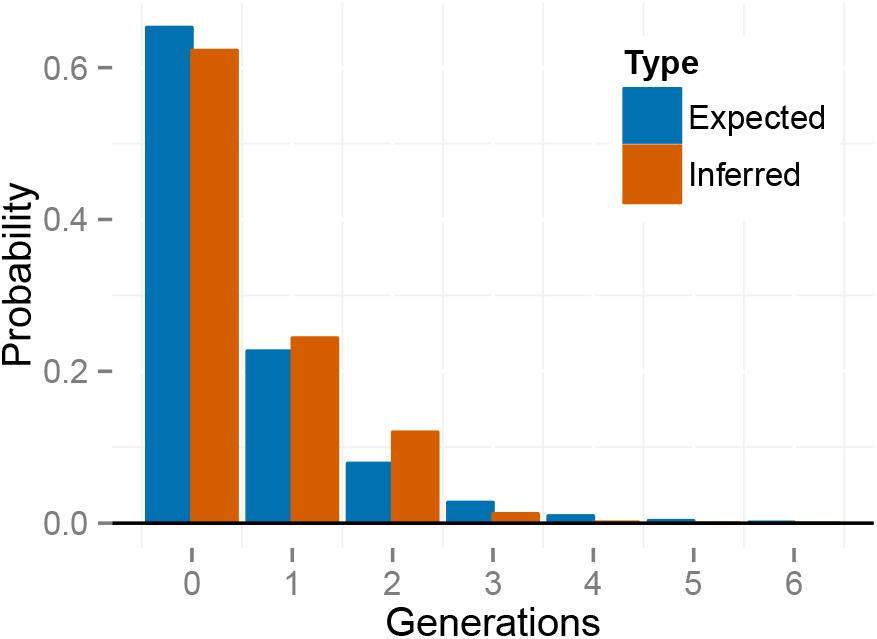
Empirical distribution of selfing times *T* across individuals, for *K. marmoratus* (Population TC). The histogram is averaged across posterior samples.

### Gynodioecious plant

We next examine data from *Schiedea salicaria*, a gynodioecious member of the carnation family endemic to the Hawaiiian islands. We analyzed genotypes at 9 microsatellite loci from 25 *S. salicaria* individuals collected from west Maui and identified by Wallace *et al.* (2011) as non-hybrids.

#### Parameter estimation

Our gynodioecy model (27) comprises 4 basic parameters, including the relative seed set of females (*σ*) and the relative viability of uniparental offspring (*τ*). Our analysis of microsatellite data from the gynodioecious Hawaiian endemic *Schiedea salicaria* (Wallace *et al.* 2011) constrained the relative seed set of females to unity (*σ ≡* 1), consistent with empirical results (Weller and Sakai 2005). In addition, we use results of experimental studies of inbreeding depression to develop an informative prior distribution for *τ*:

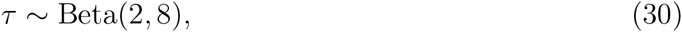

the mean of which (0.2) is consistent with the results of greenhouse experiments reported by Sakai *et al.* (1989).

Campbell *et al.* (2010) reported a 12% proportion of females (*n*_*f*_ = 27 females among *n*_*total*_ = 221 individuals). As in the case of androdioecy (29), we model this information by

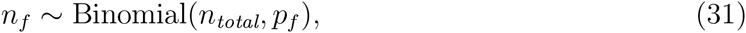

obtaining estimates from the extended likelihood function corresponding to the product of Pr(*n*_*f*_ *|n*_*total*_, *p*_*f*_) and the likelihood of the genetic data. We retain a uniform prior for the proportion of seeds of hermaphrodite set by self-pollen (*a*).

#### Results

Figure S10 (Supplementary Material) presents posterior distributions of the basic parameters of the gynodioecy model (10). Our estimate of the uniparental proportion *s*_*G*_ (median 0.247, 95% BCI (.0791, 0.444)) is substantially lower than the *F*_*IS*_-based estimate (28) of *s*_*G*_ = 0.33. Although 

~~~
RMES
~~~

 excluded none of the loci, it gives an estimate of *s*_*G*_ = 0, with a 95% CI of (0, 0.15).

Unlike the *K. marmoratus* data sets, the *S. salicaria* data set does not appear to provide substantial evidence for large differences in locus-specific mutation rates across loci: Figure S8 (Supplementary Material) shows similar posterior medians for across loci.

Figure 11 presents the inferred distribution of the number of generations since the most recent outcross event *T* across individuals, averaged over posterior uncertainty. In contrast with the analysis of the *K. marmoratus* BP population (Figure 8), the distribution appears to be consistent with the inferred uniparental proportion *s*_*G*_. Figure S9 (Supplementary Material) presents the posterior distribution of the number of consecutive generations of selfing in the immediate ancestry of each individual.

**Figure 11.**
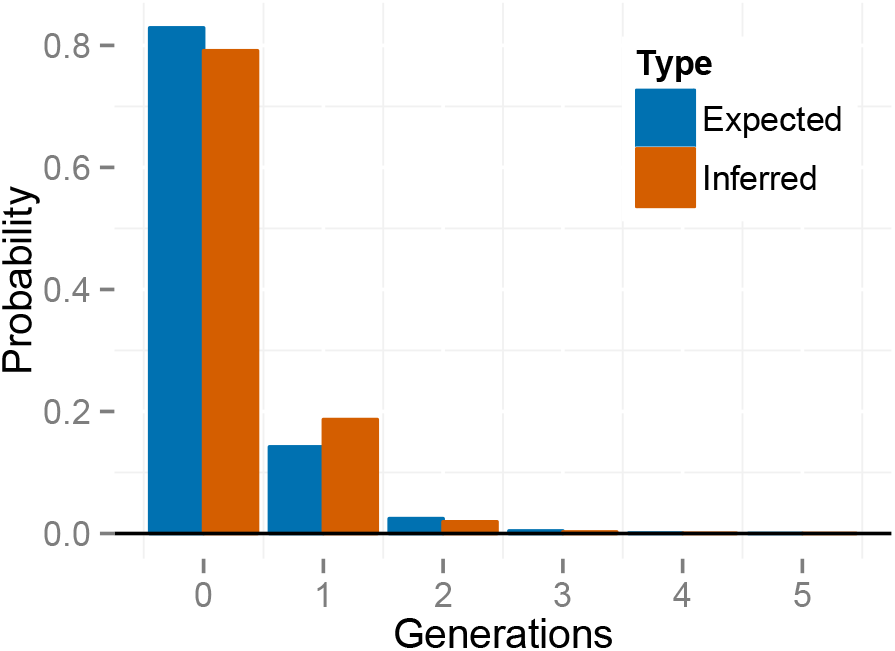
Empirical distribution of selfing times *T* across individuals, for *S. salicaria*. The histogram is averaged across posterior samples.

Table 1 presents posterior medians and 95% BCIs for the proportion of uniparentals among reproductives (*s\**), the proportion of seeds set by hermaphrodites by self-pollen (*a*), the viability of uniparental offspring relative to biparental offspring (*τ*), the proportion of females among reproductives (*p*_*f*_), and the probability that a random gene derives from a female parent ((1 *- s*_*G*_)*F/*2). Comparison of the first (YYY) and fifth (NYY) rows indicates that inclusion of the genetic data more than doubles the posterior median of *s\** (from 0.112 to 0.247) and shrinks the credible interval. Comparison of the first (YYY) and third (YNY) rows indicates that counts of females and hermaphrodites greatly reduce the posterior median of *p*_*f*_ and accordingly change the proportional contribution of females to the gene pool ((1 *- s*_*G*_)*F/*2). The bottom row of the table (NNN), showing a prior estimate for composite parameter *s\** of 0.0844 (0.000797, 0.643), illustrates that its induced prior distribution departs from uniform on (0, 1), even though both of its components (*a* and *τ*) have uniform priors.

**Table 1.** Parameter estimates for different amounts of data. Estimates are given by a posterior median and a 95% BCI.

| G | F | I | $s^*$ | $a$ | $\tau$ | $p_f$ | $(1 - s_G)F/2$ |
| --- | --- | --- | --- | --- | --- | --- | --- |
| Y | Y | Y | 0.247 (0.0791, 0.444) | 0.695 (0.299, 0.971) | 0.215 (0.0597, 0.529) | 0.125 (0.0849, 0.173) | 0.118 (0.054, 0.258) |
| Y | Y | N | 0.267 (0.0951, 0.469) | 0.497 (0.187, 0.93) | 0.507 (0.082, 0.973) | 0.125 (0.0851, 0.174) | 0.0808 (0.0398, 0.191) |
| Y | N | Y | 0.213 (0.045, 0.402) | 0.742 (0.379, 1.00) | 0.252 (0.0488, 0.529) | 0.244 (0.00, 0.613) | 0.218 (0.0, 0.403) |
| Y | N | N | 0.243 (0.0608, 0.429) | 0.628 (0.268, 0.999) | 0.611 (0.167, 1.00) | 0.354 (0.00, 0.072) | 0.223 (0.00, 0.394) |
| N | Y | Y | 0.112 (0.0026, 0.588) | 0.496 (0.0252, 0.974) | 0.183 (0.0277, 0.513) | 0.125 (0.0847, 0.173) | 0.0956 (0.0427, 0.218) |
| N | Y | N | 0.231 (0.00391, 0.776) | 0.504 (0.025, 0.973) | 0.493 (0.0257, 0.975) | 0.125 (0.0847, 0.173) | 0.0778 (0.0392, 0.172) |
| N | N | Y | 0.0376 (0.00, 0.318) | 0.492 (0.0122, 0.957) | 0.0.185 (0.00917, 0.462) | 0.483 (0.00, 0.946) | 0.314 (0.0361, 0.500) |
| N | N | N | 0.0844 (0.000, 0.643) | 0.497 (0.0244, 0.975) | 0.494 (0.0252, 0.975) | 0.479 (0.0245, 0.972) | 0.289 (0.0313, 0.5) |

## Discussion

We introduce a model-based Bayesian method for the inference of the rate of self-fertilization and other aspects of a mixed mating system. In anticipation of large (even genome-scale) numbers of loci, it uses the Ewens Sampling Formula (ESF) to determine likelihoods in a computationally efficient manner from frequency spectra of genotypes observed at multiple unlinked sites throughout the genome. Our MCMC sampler explicitly incorporates the full set of parameters for each iconic mating system considered here (pure hermaphroditism, androdioecy, and gynodioecy), permitting insight into various components of the evolutionary process, including effective population size relative to the number of reproductives.

### Assessment of the new approach

#### Accuracy

Enjalbert and David (2000) and David *et al.* (2007) base estimates of selfing rate on the distribution of numbers of heterozygous loci. Both methods strip genotype information from the data, distinguishing between only homozygotes and heterozygotes, irrespective of the alleles involved. Loci lacking heterozygotes altogether (even if polymorphic) are removed from the analysis as uninformative about the magnitude of departure from Hardy-Weinberg proportions (Figure 4). As the observation of polymorphic loci with low heterozygosity provides strong evidence of inbreeding, exclusion of such loci by 

~~~
RMES
~~~

 (David *et al.* 2007) may contribute to its loss of accuracy for high rates of selfing (Figure 2).

Our method derives information from all loci. Like most coalescence-based models, it accounts for the level of variation as well as the way in which variation is partitioned within the sample. Even a locus monomorphic within a sample provides information about the age of the most recent common ancestor of the observed sequences, a property that was not widely appreciated prior to analyses of the absence of variation in a sample of human Y chromosomes (Dorit *et al.* 1995; Fu and Li 1996).

Estimates of the rate of inbreeding produced by our method appear to show greater accuracy than 

~~~
RMES
~~~

 and the *F*_*IS*_-based method (28) over much of the parameter range (Figure 2). The increased error exhibited under very high rates of inbreeding (*s* ≈* 1) may reflect violation of our assumption (5) that random outcrossing occurs on a much shorter time scale than mutation and coalescence. Even though our method assumes that the rate of inbreeding lies in (0, 1), the posterior distribution for data generated under random outcrossing (*s\** = 0) does indicate greater confidence in low rates of inbreeding (Figure 3).

**Figure 3.**
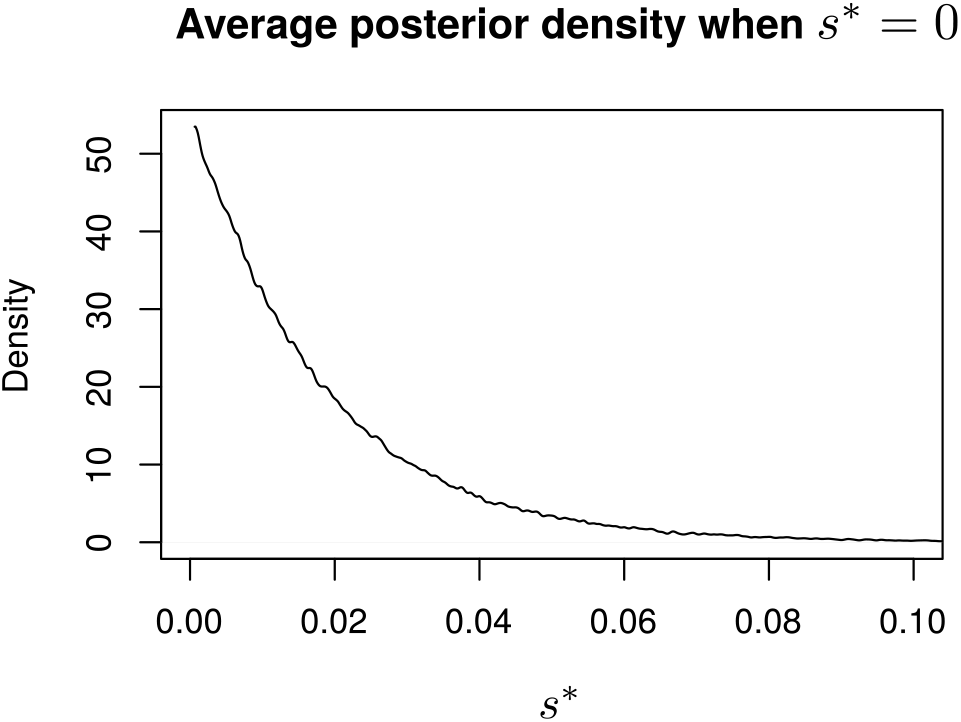
Average posterior density of the uniparental proportion (*θ\**) inferred from simulated data generated under the large sample regime (*n* = 70, *L* = 32) with a true value of *s\** = 0. The average was taken across posterior densities for 100 data sets.

Both 

~~~
RMES
~~~

 and our method invoke independence of genealogical histories of unlinked loci, conditional on the time since the most recent outcrossing event. 

~~~
RMES
~~~

 seeks to approximate the likelihood by summing over the distribution of time since the most recent outcross event, but truncates the infinite sum at 20 generations. The increased error exhibited by 

~~~
RMES
~~~

 under high rates of inbreeding may reflect that the likelihood has a substantial mass beyond the truncation point in such cases. Our method explicitly estimates the latent variable of time since the most recent outcross for each individual (14). This quantity ranges over the non-negative integers, but values assigned to individuals are explored by the MCMC according to their effects on the likelihood.

#### Frequentist coverage properties

Bayesian approaches afford a direct means of assessing confidence in parameter estimates, and our simulation studies suggest that the Bayesian Credible Intervals (BCIs) generated by our method have relatively good frequentist coverage properties as well (Figure S3). The Confidence Intervals (CIs) reported by the maximum-likelihood method 

~~~
RMES
~~~

 (David *et al.* 2007) appear to perform less well (Figure 5). Although David *et al.* (2007) describe 

~~~
RMES
~~~

 as determining CIs via the profile likelihood method (see Kreutz *et al.* 2013), 

~~~
RMES
~~~

 holds constant parameters other than the uniparental proportion (*s\**) instead of reoptimizing them to maximize the likelihood as *s\** varies. The result is therefore not a true profile likelihood, which may explain the poor coverage properties of the CIs that 

~~~
RMES
~~~

 provides.

#### Model fit

Bayesian approaches also afford insight into the suitability of the underlying model. Our method provides estimates of the number of generations since the most recent outcross event in the immediate ancestry of each individual (*T*). We can pool such estimates of selfing times to obtain an empirical distribution of the number of selfing generations, a procedure particularly useful for samples containing observation of the genotype of many individuals. Under the assumption of a single population-wide rate of self-fertilization, we expect selfing time to have a geometric distribution with parameter corresponding to the estimated selfing rate. Empirical distributions of the estimated number of generations since the last outcross appear consistent with this expectation for the data sets derived from the TC population of *K. marmoratus* (Figure 10) and from *Schiedea* (Figure 11). In contrast, the empirical distribution for the highly-inbred BP population of *K. marmoratus* (Figure 8) shows an absence of individuals formed by random outcrossing (*T* = 0). That our method accurately estimates *T* from simulated data (Figure 6) argues against attributing the inferred deficiency of biparental individuals in the BP data set to an artifact of the method. Rather, the deficiency may indicate a departure from the underlying model, which assumes reproduction only through self-fertilization or random outcrossing. In particular, biparental inbreeding as well as selfing may reduce the fraction of individuals formed by random out-crossing. Mis-scoring of heterozygotes as homozygotes due to null alleles or other factors, a possibility directly addressed by 

~~~
RMES
~~~

 (David *et al.* 2007), may also in principle contribute to the paucity of outbred individuals.

### Components of inference

#### Locus-specific mutation rates

Our method estimates the scaled mutation rate (4) at each locus using the Dirichlet Process Prior (DPP). This approach improves on existing methods in several ways. First, we estimate a single parameter for each locus instead of estimating multiple allele frequencies per locus as do Enjalbert and David (2000). Second, we estimate for each locus the scaled mutation rate, a fundamental component of the evolutionary process, rather than the heterozygosity (1), a random outcome of that process. Third, incorporation of the DPP permits the simultaneous estimation of the number of classes of mutation rates, the mutation rate for each class, and the class membership of each locus. It accords the increased accuracy derived from pooling loci with similar mutation rates without *a priori* knowledge of the partitioning of loci among rate classes or even the number of classes.

#### Joint inference of mutation and inbreeding rates

For the infinite-alleles model of mutation, the Ewens Sampling Formula (ESF, Ewens 1972) provides the probability of any allele frequency spectrum (AFS) observed at a locus in a sample derived from a panmictic population. Under partial self-fertilization, the ESF provides the probability of an AFS observed among genes, each sampled from a distinct individual. For such genic (as opposed to genotypic) samples, the coalescence process under inbreeding is identical to the standard coalescence process, but with a rescaling of time (Fu 1997; Nordborg and Donnelly 1997). Accordingly, genic samples may serve as the basis for the estimation of the single parameter of the ESF, the scaled mutation rate *θ\** (6), but not the rate of inbreeding apart from the scaled mutation rate.

Our method uses the information in a genotypic sample, the genotype frequency spectrum, to infer both the uniparental proportion *s\** and the scaled mutation rate *θ\**. Our sampler reconstructs the genealogical history of a sample of diploid genotypes only to the point of the most recent random-outcross event of each individual, with the number of consecutive generations of inbreeding in the immediate ancestry of a given individual (*T*_*k*_ for individual *k*) corresponding to a latent variable in our Bayesian inference framework. Invocation of the ESF beyond the point at which all lineages reside in separate individuals obviates the necessity of further genealogical reconstruction. As a consequence, our method may be better able to accommodate genome-scale magnitudes of observed loci (*L*).

Identity disequilibrium (Cockerham and Weir 1968), the correlation in heterozygosity across loci within individuals, reflects that all loci within an individual experience the most recent random-outcross event at the same time, irrespective of physical linkage. The heterozygosity profile of individual *k* provides information about *T*_*k*_ (16), which in turn reflects the uniparental proportion *θ\**. Observation of multiple individuals provides a basis for inference of both the uniparental proportion *s\** and the scaled mutation rate *θ\**.

#### Identifiability

In an analysis based solely on the genotype frequency spectrum observed in a sample, the likelihood depends on just two composite parameters: the probability that a random individual is uniparental (*s\**) and the scaled rates of mutation **Θ\*** (20) across loci. Any model for which the parameter set **Ψ** (24) comprises more than one parameter is not fully identifiable from the genetic data alone. In the pure hermaphroditism model (7), for example, basic parameters 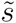 (fraction of fertilizations by selfing) and *τ* (relative viability of uniparental offspring) are nonidentifiable: any assignments that determine the same values of composite parameters *s\** and **Θ\*** have the same likelihood.

For each basic parameter in **Ψ** beyond one, identifiability requires incorporation of additional information beyond the genetic data. A full treatment of such information requires expansion of the likelihood function to encompasses an explicit model of the new information. Our androdioecy model (8), for example, comprises 3 parameters, including the frequency of males among reproductives (*p*_*m*_) as well as 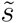 and *τ*. In our analysis of microsatellite data from the killifish *Kryptolebias marmoratus* (Mackiewicz *et al.* 2006; Tatarenkov *et al.* 2012), the expanded likelihood function corresponds to the product of the probability of the genetic data and the probability of the number of males observed among a total number of individuals (29). In the absence of information regarding inbreeding depression (*τ*), we assigned *τ ≡* 1 to permit estimation of the uniparental proportion (*s\**) under a uniform prior distribution. This assignment does not affect the reliability of our estimates (*s\**, **Θ\***, *p*_*m*_, *S*_*A*_, *etc*.); rather, the analysis is agnostic concerning the influence of the relative viability of inbred offspring (*τ*) and the rate of self-fertilization 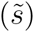 in determining the probability that a random individual is uniparental (s*\**).

Non-identifiable parameters can also be estimated through the incorporation of informative priors. Because identifiability is defined in terms of the likelihood, which is unaffected by priors, such parameters remain non-identifiable. Even so, informative priors assist in their estimation through Bayesian approaches, which do not require parameters to be identifiable. To explore the data set from *Schiedea salicaria* (Wallace *et al.* 2011), we use our 4-parameter gynodioecy model (10), the basic parameters of which include the proportion of females among reproductives (*p*_*f*_), the relative seed set of females (*σ*), the relative viability of uniparental offspring (*τ*), and the proportion of seeds of hermaphrodites set by self-pollen (*a*). In a manner similar to the androdioecy study, our analysis uses an extended likelihood function, modeling the number of females as a binomial random variable (31). In addition, we use earlier experimental evidence to justify the assignment of *σ* ≡ 1 (Weller and Sakai 2005) and to develop an informative prior for *τ* ((30): Sakai *et al.* 1989). This procedure permits estimation of 3 basic parameters, including the proportion of seeds of hermaphrodites set by self-pollen (*a*).

### Beyond estimation of the selfing rate

Our MCMC implementation updates the full set of basic parameters, with likelihoods determined from the implied values of composite parameters *s\** and **Θ\***. Incorporation of additional information, either through extension of the likelihood or through informative priors, permits inference not only of the basic parameters but also of functions of the basic parameters. For example, Table 1 includes estimates of the proportion of seeds of hermaphrodites set by self-pollen (*a*) and the probability that a random gene derives from a female parent ((1*-s*_*G*_)*F/*2) in gynodioecious *S. salicaria*. We are not aware of other studies in which these quantities have been inferred from the pattern of neutral genetic variation observed in a random sample.

Among the most biologically-significant functions to which this approach affords access is relative effective number *S* (4), a fundamental component of the reproductive value of the sexes (Fisher 1958). We denote the probability that a pair of genes, randomly drawn from distinct individuals, derive from the same parent in the preceding generation as the rate of parent-sharing (1*/N \**). Its inverse (*N \**) corresponds to the “inbreeding effective size” of Crow and Denniston (1988). Relative effective number *S* is the ratio of *N \** to the total number of reproductive individuals. For example, in the absence of inbreeding (*s\** = 0), *N \** in our gynodioecy model (10) corresponds to Wright’s 1969 harmonic mean expression for effective population size and *S* to the ratio of *N \** and *N*_*f*_ + *N*_*h*_, the total number of reproductive females and hermaphrodites. In general (*s* ≥* 0), relative effective size *S* reflects reductions in effective size due to inbreeding in addition to differences in numbers of the sexual forms. Figure 12 presents posterior distributions of *S* for the 3 data sets explored here. These results suggest that relative effective number *S* in each of the natural populations surveyed lies close to its maximum of unity, with the effective number defined through the rate of parent-sharing approaching the total number of reproductives.

**Figure 12.**
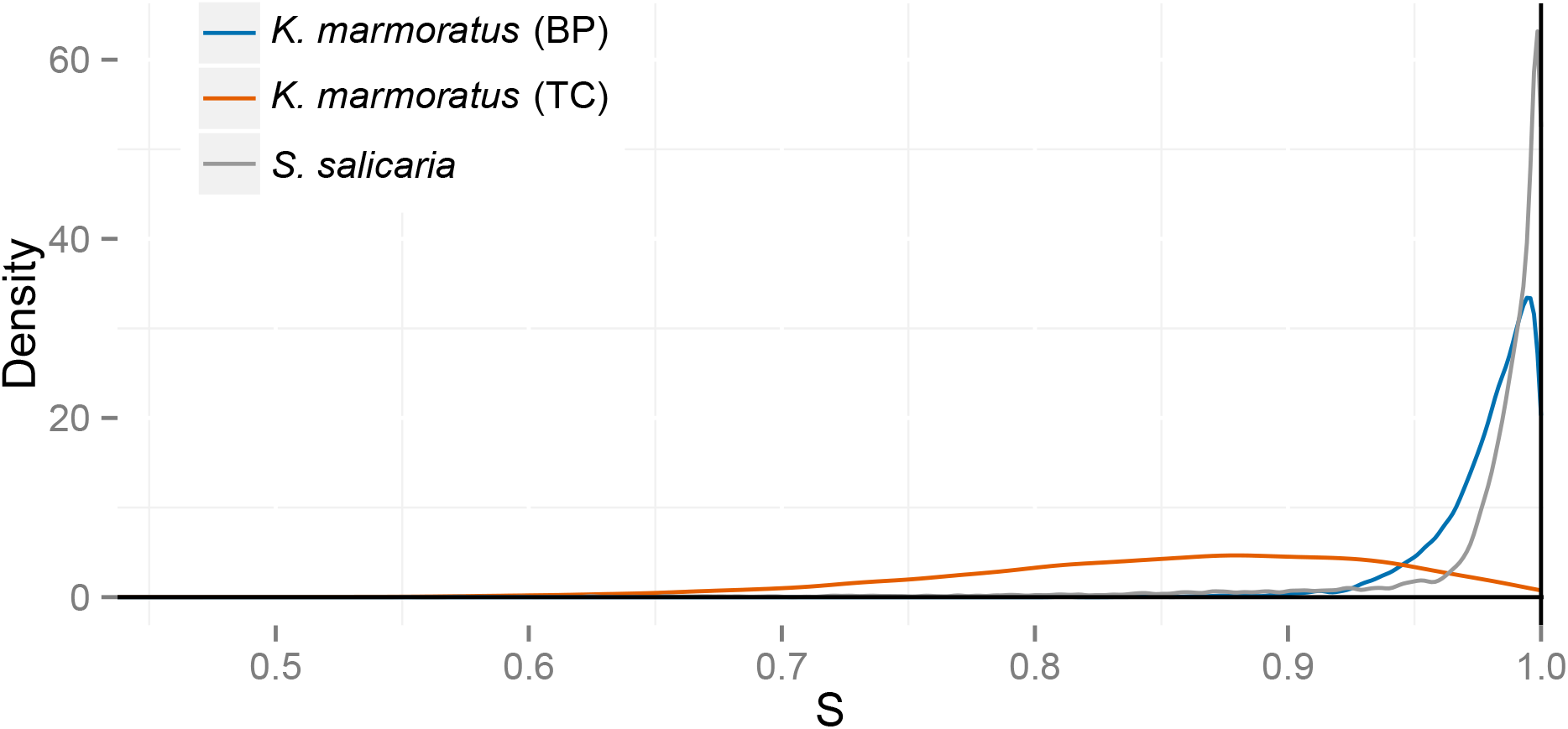
Posterior distributions of relative effective number *S* (4) for data sets derived from *Kryptolebias marmoratus* (BP and TC populations) and *Schiedea salicaria*.

## ACKNOWLEDGMENTS

This project was initiated during a sabbatical visit of MKU to the Department of Ecology and Evolutionary Biology at the University of California at Irvine. We thank Francisco J. Ayala for exceedingly gracious hospitality, and Diane R. Campbell and all members of the Department for stimulating interactions. We thank Lisa E. Wallace for making available to us microsatellite data from *Schiedea salicaria*. Public Health Service grant GM 37841 (MKU) provided partial funding for this research.

## Appendix A The last-sampled gene

We address the probability that the last-sampled gene in a sample of size *i* represents a novel allele (22a).

Under the infinite alleles model of mutation, a single mutation in a lineage suffices to distinguish a new allele. We denote the last-sampled gene in a sample of size *i* as the focal gene, and consider the level of the genealogical tree in which its ancestral lineage either receives a mutation or joins the gene tree of the sample at size (*i -* 1). Level *l* of the entire (*i*-gene) gene tree corresponds to the segment in which *l* lineages persist.

The probability that the line of descent of the focal gene terminates in a mutation immediately, in level *i* of the genealogy, is

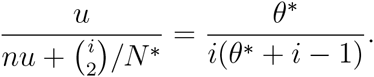

In general, the probability that the lineage of the focal gene terminates on level *l >* 2 is

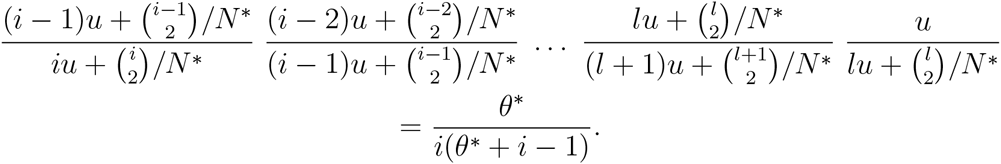

This expression illustrates the invariance over termination orders noted by Griffiths and Lessard (2005). Summing over all levels, including level 2, for which a mutation in either remaining lineage ensures that the focal gene represents a novel allele, we obtain the overall probability that the last-sampled gene represents a novel allele:

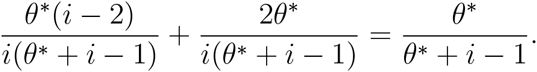

## Appendix B Estimators of *F* _*IS*_

We follow Weir (1996) in developing an estimate of the uniparental proportion *s\** from *F*_*IS*_ alone (28).

For a single locus, a simple estimator of *F*_*IS*_ corresponds to

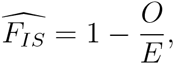

for *O* the observed fraction of heterozygotes in the sample and *E* the expected fraction based on Hardy-Weinberg proportions given the observed allele frequencies. Explicitly, we have

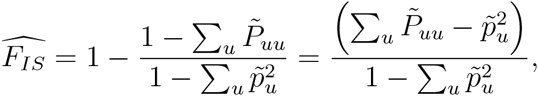

for 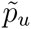 the frequency of allele *u* in the sample and 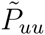 the frequency of homozygous genotype *uu* in the sample. However, this estimator can be substantially biased for small samples, leading to underestimation of *F*_*IS*_ (Weir 1996).

To address this bias and accommodate multiple loci, we instead adopt

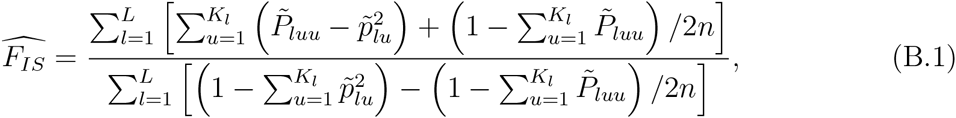

for *n* the number of diploid genotypes observed, *L* the number of loci, and *K*_*l*_ the number of alleles at locus *l*. While this estimator is also biased in general, it corresponds to the ratio of unbiased estimators of 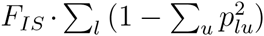 and 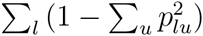, in which *p*_*lu*_ is the frequency of allele *u* at locus *l* in the entire population (Weir 1996). Our analysis of simulated data (Appendix D) indicates that this estimator is more accurate than an estimator that simply averages single-locus estimates:

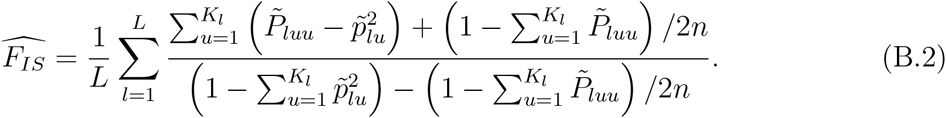

Our *F*_*IS*_-based estimates (28) incorporate (B.1) and not (B.2).

## Appendix C Implementation of the MCMC

### State space

The state space for the Markov chain of our MCMC sampler includes times across sampled individuals since the last outcross event **T** (14), coalescence events across individuals and loci since that event **I** (15), and model-specific parameters **Ψ** (24). The state space also comprises the scaled mutation rates **Θ\*** (20), which are determined by **C**, a list specifying the mutation rate category *C*_*l*_ for locus *l* = 1 *… L*, and **Z**, a list specifying the scaled mutation rate *Z*_*i*_ for category *i* = 1 *… L* + 4. In particular, the scaled mutation rate at locus *l* corresponds to

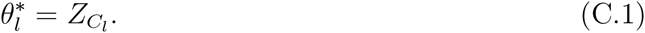

At any given point in the MCMC, the state of the Markov chain corresponds to (**I**, **T**, **Ψ**, **C**, **Z**).

### Iterations

Each iteration of our MCMC sampler performs multiple updates, with each variable updated at least once per iteration. We recorded the state sampled by the MCMC at each iteration. For analyses of simulated data sets, we ran Markov chains for 2000 iterations, discarding the first 200 iterations as burn-in. For analyses of the actual data sets, we ran Markov chains for 100,000 iterations, discarding the first 10,000 iterations as burn-in. Convergence appeared to occur as rapidly for actual data as for simulated data, but we found empirically that the larger number of samples were needed to achieve smooth density plots for the actual data sets.

### Transition kernels

Updating of the continuous variables of mutation rates *{Z*_*l*_*}* (C.1) and model-specific parameters **Ψ** (24) uses both Metropolis-Hastings (MH) transition kernels and auto-tuned slice-sampling transition kernels. Updating of the discrete variables *{C*_*l*_*}* uses a Gibbs transition kernel.

### Efficient inference on selfing times through collapsed Metropolis-Hastings

Simple Metropolis-Hastings (MH) proposals that separately update the time since the most recent outcross event (*T*_*k*_) and coalescence history since that event (*I*_*·k*_) lead to extremely poor mixing efficiency. Strong correlations between *T*_*k*_ and *I*_*·k*_ cause changes to *T*_*k*_ to be rejected with high probability unless *I*_*·k*_ is updated as well. For example, consider proposing a change of *T*_*k*_ from 1 to 0. When *T*_*k*_ = 1, on average *I*_*lk*_ will be 1 at half of the loci and 0 at the remaining loci. If any of the *I*_*lk*_ = 1, a move to *T*_*k*_ = 0 will always be rejected because the probability of a coalescence event more recently than the most recent outcross event is 0 if the sampled individual is itself a product of outcrossing. To permit acceptance of changes to *T*_*k*_, we introduce a proposal for *T*_*k*_ that also changes *I*_*·k*_.

The scheme starts from the value *T*_*k*_ = *t*_*k*_ and proposes a new value 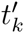. In standard MH within Gibbs, we would compute the probability of *T*_*k*_ = *t*_*k*_ and of *T*_*k*_ = *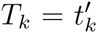* given that all other parameters are unchanged. We modify this MH scheme to compute probabilities without conditioning on the coalescence indicators for individual *k*. However, the coalescence indicators for other individuals are still held constant. To compute this probability, let *J* indicate all the coalescence indicators *I*_*·y*_ where *y ≠ k*. Then

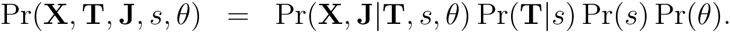

We introduce **I**_*·k*_ by summing over all possible values **i**_*·k*_.

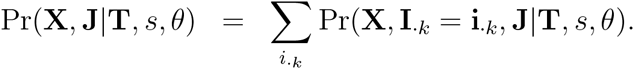

Since the *i*_*lk*_ for different loci are independent given *T*_*k*_, we have

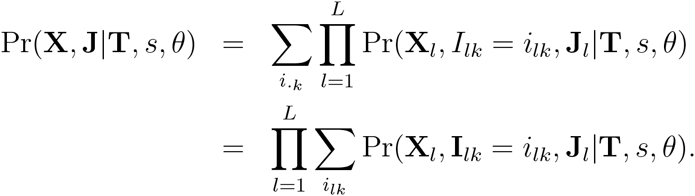

Therefore, for specific values of **T** and **J**, we can compute the sum over all possible values of **I**_*·k*_for *l* = 1 *… L* in computation time proportional to *L* instead of 2^*L*^. This is possible because the *L* coalescence indicators for individual *k* each affect different loci, and are conditionally independent given *T*_*k*_ and **J**.

After accepting or rejecting the new value of *T*_*k*_ with *I*_*·k*_ integrated out, we must choose new values for **I**_*·k*_ given the chosen value of *T*_*k*_. Because of their conditional independence, we may separately sample each coalescence indicator *I*_*lk*_ for *l* = 1 *… L* from its full conditional given the chosen value of *T*_*k*_. This completes the collapsed MH proposal.

## Appendix D Analysis of simulated data

**Simulations:** Our simulator (

~~~
https://github.com/skumagai/selfingsim
~~~

) was developed using 

~~~
simuPOP
~~~

, publicly available at 

~~~
http://simupop.sourceforge.net/
~~~

. It explicitly represents *N* = 10, 000 individuals, each bearing two genes at each of *L* unlinked loci. Mutations arise at locus *l* at scaled rate *θ*_*l*_ (4), in accordance with the the infinite-alleles model.

We assigned to uniparental proportion *s\** values ranging from 0.01 to 0.99, with half of the *L* = 32 loci assigned scaled mutation rate *θ* = 0.5 and the remaining loci *θ* = 1.5.

We conducted 10^2^ independent simulations for each assignment of *s\**. Each simulation was initialized with each of the 2*N ×* 32 genes representing a unique allele. Most of this maximal heterozygosity was lost very rapidly, with allele number and allele frequency spectrum typically stabilizing well within 10*N* generations. After 20*N* generations, we recorded the realized population, from which 100 independent samples of *L* = 32 loci of size *n* = 70 were extracted. From this collection, we randomly chose *L* = 6 loci and subsampled 100 independent samples of size *n* = 6.

### Analysis

To 10^2^ independent samples from each of 10^2^ independent simulations for each assignment of the uniparental proportion *θ\**, we applied our Bayesian method, the *F*_*IS*_ method, and 

~~~
RMES
~~~

. Our Bayesian method is open-source and can be obtained at

https://github.com/bredelings/BayesianEstimatorSelfing/.

We used the implementation of 

~~~
RMES
~~~

 (David *et al.* 2007) provided at

http://www.cefe.cnrs.fr/images/stories/DPTEEvolution/Genetique/fichiers% 20Equipe/RMES%202009%282%29.zip.

